# Resolution of conflict between parental genomes in a hybrid species

**DOI:** 10.1101/102970

**Authors:** Fabrice Eroukhmanoff, Richard I. Bailey, Tore O. Elgvin, Jo S. Hermansen, Anna R. Runemark, Cassandra N. Trier, Glenn-Peter Sætre

## Abstract

The development of reproductive barriers against parent species is crucial during hybrid speciation, and post-zygotic isolation can be important in this process. Genetic incompatibilities that normally isolate the parent species can become sorted in hybrids to form reproductive barriers towards either parent. However, the extent to which this sorting process is systematically biased and therefore predictable in which loci are involved and which alleles are favored is largely unknown. Theoretically, reduced fitness in hybrids due to the mixing of differentiated genomes can be resolved through rapid evolution towards allelic combinations ancestral to lineage-splitting of the parent species, as these alleles have successfully coexisted in the past. However, for each locus, this effect may be influenced by its chromosomal location, function, and interactions with other loci. We use the Italian sparrow, a homoploid hybrid species that has developed post-zygotic barriers against its parent species, to investigate this prediction. We show significant bias towards fixation of the ancestral allele among 57 nuclear intragenic SNPs, particularly those with a mitochondrial function whose ancestral allele came from the same parent species as the mitochondria. Consistent with increased pleiotropy leading to stronger fitness effects, genes with more protein-protein interactions were more biased in favor of the ancestral allele. Furthermore, the number of protein-protein interactions was especially low among candidate incompatibilities still segregating within Italian sparrows, suggesting that low pleiotropy allows steep intraspecific clines in allele frequencies to form. Finally, we report evidence for pervasive epistatic interactions within one Italian sparrow population, particularly involving loci isolating the two parent species but not hybrid and parent. However there was a lack of classic incompatibilities and no admixture linkage disequilibrium. This suggests that parental genome admixture can continue to constrain evolution and prevent genome stabilization long after incompatibilities have been purged.

## INTRODUCTION

The role of epistasis (non-additive interactions between alleles at different loci) in the evolution of reproductive isolation has been extensively investigated in the past years (Coyne and Orr 2004). Speciation often results from the accumulation of *de novo* mutations (Dobzhansky 1940) that, while having no deleterious effect within species, can yield allelic combinations in hybrids that make them inviable or sterile. Both theoretical models (Bateson 1909; Dobzhansky 1937; Muller 1939) and empirical evidence confirm that such Dobzhansky-Muller incompatibilities (DMIs) are important in speciation (Coyne & Orr 2004; Cutter 2012; Crespi and Nosil 2013; Seehausen et al. 2014). Yet, hybridization is widespread (Mallet 2005) and has a variety of evolutionary consequences, including the formation of hybrid species (Mallet 2007; Abbott et al. 2013). How DMIs shape hybrid genome evolution is, however, still poorly understood.

One way in which hybrid populations can escape outbreeding depression is through rapid purging of DMIs (see e.g. Rieseberg 1997; Buerkle & Rieseberg 2008). There is for instance experimental support for selection favoring the retention of genotypes that interact favorably (Rieseberg et al. 1996; Rieseberg 1997; Eroukhmanoff et al. 2013a) thus leading to rapid fixation of such genotypes. Ancestral genotypes represent viable allelic combinations that were present in the ancestral species, prior to the lineage-splitting event that resulted in the two currently hybridizing species. Ancestral alleles are thus intrinsically less likely to be involved in detrimental epistatic interactions than derived alleles (*de novo* mutations having occurred within either of the two parent species after lineage-splitting (Orr 1995; Sherman et al. 2014)). Hence, fixation of alleles at incompatibility loci in hybrid populations is expected to be highly asymmetric in favor of recreating the ancestral allelic state, reducing incompatibility with other loci present in the admixed genome (Orr 1995; Gavrilets 1997). The hypothesis that reconstruction of fit ancestral allele combinations occurs in hybrid populations has been discussed in studies of fitness consequences of DMIs (Shuker et al. 2005; Sherman et al. 2014), but has yet to be formally tested in any ancient admixed population, let alone in a hybrid species.

If DMIs involve interactions between the nuclear genome and a non-recombining organellar genome such as that of mitochondria, which may contain many strongly differentiated loci in linkage disequilibrium, this can impose stronger selection on nuclear genes towards reconstruction of the ancestral state. For instance, we should expect that nuclear genes involved in metabolic functions (e.g. genes involved in the OXPHOS system (Zhang and Broughton 2013)) are likely to interact with mitochondria and therefore cause incompatibilities when hybridization occurs (Burton and Barreto 2012). In hybrid lineages, a bias towards fixation of ancestral alleles specifically at loci where the ancestral allele is from the same parent species as their mitochondria may then occur. Derived alleles from the same parent should be compatible with the mitochondria, and hence not selected against. Purging may also be stronger and more ancestry-biased on sex chromosomes compared to autosomes, as genes involved in reproductive isolation are expected to be more common and more exposed to selection on sex chromosomes (Charlesworth et al. 1987; Qvarnström and Bailey 2009).

The persistence of DMIs in hybrids could have important consequences for the evolutionary potential of hybrid populations, either enhancing or interfering with other forms of selection (Bailey et al. 2013). Recently, an empirical study demonstrated that there can be great variation in DMI genetic architecture (Sherman et al. 2014), and this may influence their persistence. Incompatibility loci involved in simple pairwise epistatic interactions should rapidly evolve towards fit allele combinations (Orr 1995; Coyne and Orr 2004). However, more complex epistasis, which is expected for highly pleiotropic loci, may alter selective pressures. On the one hand, increasing complexity of interactions increases the likelihood of a locus interacting with derived alleles from both parent species, potentially constraining purging by causing greater antagonistic selection and reduced directional selection acting to purge unfit alleles (Rieseberg 1997; Sherman et al. 2014). Such high pleiotropy may in general prevent a locus from evolving independently of the rest of the genome, and is known to slow down evolution (Mank et al. 2008; Papakostas et al. 2014; Uebbing et al. 2016). On the other hand, a higher number of interacting loci may lead to stronger total selection favoring the ancestral allele, making purging more effective. Consequently, high pleiotropy could increase the likelihood of a moving allele frequency cline overcoming geographic and demographic barriers as it spreads in space (Barton and Hewitt 1985). High pleiotropy may therefore either improve purging if it increases selection favoring the ancestral allele, or alternatively lead to antagonistic and hence weaker selection so that moving clines becoming trapped by population density troughs, or physical or environmental barriers (Barton and Hewitt 1985; Mallet 1993; Barton and de Cara 2010; Bierne et al. 2011).

To test whether incompatibilities mold hybrid genome evolution and canalize genomes towards reconstruction of ancestral genotypes, one should ideally investigate admixed genomes that have had time to evolve in isolation from parental gene flow. Homoploid hybrid species provide such study systems, as by definition, gene flow from the parental species is restricted or absent. Homoploid hybrid speciation (HHS) is the process through which interbreeding between two taxa results in a third, novel taxon with the same number of chromosome sets, which remains distinct by means of pre-and/or post-zygotic reproductive barriers against both parent taxa (Mallet 2007; Abbott et al. 2013). In recent years, a number of putative examples of HHS have been proposed in different animal taxa (Nolte et al. 2005; Schwartz et al. 2005; Mavárez et al. 2006; Gompert et al. 2006; Elgvin et al. 2011; Hermansen et al. 2011; Gompert et al. 2014). Some of these studies present evidence that novel reproductive barriers have arisen from hybridization, but so far, postzygotic isolation through sorting of existing parental genetic incompatibilities has only been specifically tested and shown to be a key-ingredient for HHS in the Italian sparrow (*Passer italiae*) (Hermansen et al. 2014).

The Italian sparrow is a homoploid hybrid species formed through hybridization between the house sparrow (*P. domesticus*) and the Spanish sparrow (*P. hispaniolensis*; Hermansen et al. 2011; Elgvin et al. 2011). It has a broad geographic range, occupying the whole Italian peninsula and several large Mediterranean islands, and is mostly allopatric from its parents and hence free to evolve independently. However, contact zones exist with both parent species: with Spanish sparrows in the Gargano peninsula in southeast Italy and with house sparrows in a narrow hybrid zone in the Alps (Summers-Smith 1988, Hermansen et al. 2011; Trier et al. 2014; Bailey et al. 2015). Yet, the Italian sparrow constitutes a distinct taxon (Sangster et al. 2015) and shows several forms of reproductive isolation from its parents (Trier et al. 2014; Bailey et al. 2015). As in other hybrid species (see e.g. Rieseberg et al. 1995; Rieseberg et al. 1997; Gompert et al. 2014) the Italian sparrow nuclear genome is mosaic; however it has inherited its mitochondria (mtDNA) from house sparrows. At some nuclear loci, it is fixed for the house sparrow or the Spanish sparrow allele and yet at other loci, alleles from both parents are segregating (Hermansen et al. 2011; Elgvin et al. 2011; Trier et al. 2014). Recent work has shown that this mosaicism extends to genetic incompatibilities in this hybrid species. Sex-linked and mito-nuclear incompatibilities that normally isolate the parent species-possibly via the production of infertile offspring (Eroukhmanoff et al. 2016)-have been sorted in the Italian sparrow to form reproductive barriers against one or other parent species at the hybrid-parent range boundaries, while other loci appear to represent incompatibilities still segregating within the Italian sparrow’s range (Trier et al. 2014; Hermansen et al. 2014).

In this study, we investigate the role of incompatibilities in constraining and molding hybrid genome evolution using a set of primarily exonic intra-genic loci that are divergent between the parent species. If DMIs are affecting the evolution of the Italian sparrow genome we expect that: 1) parental contribution shows a general bias towards the ancestral allele; 2) this ancestry-biased purging is more stringent for nuclear loci with a mitochondrial function, specifically loci whose ancestral allele came from the same parent species as the mitochondrial genome (in this case from house sparrows); 3) increasing protein-protein interaction complexity either increases antagonistic pleiotropy, constraining directional evolution and reducing purging, or alternatively, strengthens purging by strengthening directional selection in favor of the ancestral allele; 4) incompatibilities are still segregating within Italian sparrow populations, and can be detected through deviations from Hardy-Weinberg and linkage equilibrium. We find support for the first two predictions, and also show that loci more strongly purged in favor of the ancestral allele have more complex interaction networks, while candidate unpurged intra-Italian sparrow incompatibilities have exceptionally simple interaction networks. Finally, we find strongly suggestive evidence for continuing epistatic selection acting on segregating loci within one central-Italian sparrow population.

## METHODS

### Sampling and Genotyping

We caught 89 male Italian sparrows from one population (Lago Salso: 41.5403 N; 15.8906 E) in spring 2012 using mist nets. We sampled blood from the brachial vein, and stored the blood in Queen’s lysis buffer. Authorization to catch and sample birds was obtained from the national authorities of Italy and the regional authorities of Puglia. In addition, we also sampled 15 individuals from the more distantly related tree sparrow (*P. montanus*; 10 individuals in the Gargano peninsula and 5 in the Alps) for outgroup comparisons. Authorization for sampling in the Alps, also in 2012, was obtained from appropriate Swiss and Italian authorities (see Bailey et al. 2015).

DNA was extracted using Qiagen DNeasy 96 Blood and Tissue Kits (Qiagen N.V., Venlo, Netherlands) according to the manufacturer’s instructions with the minor adjustment of adding 100μl of blood stored in buffer in the initial step. Each individual was genotyped for a set of 80 parent species-informative single nucleotide polymorphism (SNP) markers using the Sequenom MassARRAY platform at CIGENE, Norwegian University of Life Sciences, Ås, Norway. These SNPs were previously identified using transcriptome sequencing of the two parent species, house and Spanish sparrow. Hence, they are located in protein coding regions and may therefore represent biologically significant mutations (Trier et al. 2014; Hermansen et al. 2014), except for one additional SNP (CHD1Z), which is located in an intron (Elgvin et al. 2011). In addition, we included existing data from genotypes of 385 individual male Italian sparrows from 59 populations spread across mainland Italy, the Alps (including the house-Italian sparrow hybrid zone) and Sicily, as well as Spanish sparrows from Sardinia (Hermansen, et al. 2014, Trier et al. 2014). Additional information on transcriptome mining for SNPs and genotyping procedures can be found in Hermansen et al. (2014) and Trier et al. (2014).

The four sparrow species studied here all share a common ancestor, with tree sparrows having split from the common ancestor of house and Spanish sparrows approximately 6 Mya (Allende et al. 2001). Alleles shared among several closely related taxa are most likely to represent the ancestral state, present in this common ancestor, while alleles occurring in only one branch of the phylogenetic tree are more likely to be derived. When either house or Spanish sparrows share an allele with the tree sparrow this is thus likely to reflect the allele present in the most recent common ancestor of house and Spanish sparrows. Hence, sampled tree sparrow genotypes were used to assess which allele at each locus was most likely to have arisen prior to the house/Spanish sparrow split (the ancestral allele) and which was derived. We chose to focus on a subset of 57 of the parent species-informative SNPs described above, located on 15 different chromosomes (Table S1). We selected loci with >99% genotyping success and with no more than two alleles segregating across all species. We also only retained loci with a frequency of homozygotes for the minor allele <5% in the tree sparrow and also in at least one of the two parents, thus making it plausible that derived alleles may be incompatible in an admixed genome. Most of these loci were fixed in tree sparrows, with only one (*HECTD1*) having a minor allele frequency higher than 0.1 (Table S1).

Previously (Trier et al. 2014, Hermansen et al. 2014), a Bayesian genomic cline approach (Gompert and Buerkle 2011) was used to identify loci exhibiting reduced introgression or strong bias in favor of one or other parental allele compared with average genome-wide admixture (hybrid index), suggesting an association with reproductive isolation. Genomic cline analysis involves estimation of two parameters for each locus (Gompert and Buerkle 2011). The first is α, which represents the locus-specific deviation in the probability of alleles in the test population being from one or other parent species, relative to the global hybrid index, with 0 indicating no deviation from global expectation. This is analogous to cline center in geographic cline analysis. Large positive or negative estimates of α hence suggest the purging of incompatible alleles. The second is the rate parameter β, which represents the rate of transition from one allele to the other relative to changing hybrid index, and is hence analogous to cline steepness and indicates selection for or against introgression into the foreign genomic background. In our study, candidate hybrid-parent incompatibilities were significant for both parameters (significantly restricted introgression for β), while candidate intraspecific incompatibilities were always significant for β (Trier et al. 2014). There is much geographic variation in the genomic composition of this hybrid species (Hermansen et al. 2011; Eroukhmanoff et al. 2013b; Trier et al. 2014), with a broad cline in hybrid index running north-south through the Italian peninsula. Our genomic cline analyses involved estimating a single value of each of α and β for each locus using samples covering the extent of the Italian sparrow’s mainland geographic range and beyond, starting from house sparrows adjacent to the Alpine house-Italian sparrow hybrid zone, down through Italy into Sicily, and also populations of Spanish sparrows on the nearby island of Sardinia.

A total of six of the 57 loci were identified as candidates for post-zygotic isolation both between the hybrid Italian sparrow and one of its parents, through having steep genomic clines centered on one of the two hybrid-parent range boundaries (Table S1; Trier et al. 2014), and between the parent species themselves (Hermansen et al. 2014). Another six loci have been identified as candidate intraspecific incompatibility loci, i.e. incompatibility loci with steep genomic clines centered within the geographic range of phenotypically Italian sparrows, rather than at the hybrid-parent boundary (Table S1; Trier et al. 2014, Hermansen et al. 2014). All but one of the candidate loci for intrinsic isolation between hybrid and parent were sex-linked or mitochondrial (we included one mitochondrial SNP, within the ND2 gene, in the earlier study), and mito-nuclear incompatibilities were specifically isolating Italian and Spanish sparrows (Trier et al. 2014). Internal incompatibility loci (i.e. with steep clines but α closer to zero, and with the primary cline occurring within the geographic range of Italian sparrows) were more often located on autosomes. More information about each of these SNPs can be found in Supplementary Table S1.

Genomic cline analysis is prone to false positives because drift and stochasticity can also lead to steep or shifted clines (Fitzpatrick 2013). However, false negatives are less likely, and given the large geographic region over which these loci had to spread, they represent strong candidate DMIs awaiting further verification.

### Testing the ancestral genotype reconstruction hypothesis

If ancestral genotype recovery is a major mechanism during hybrid genome stabilization, and our 57 SNPs include loci that are genuinely under selection, there should be an overall bias in genomic cline α values in favor of the ancestral allele. We tested this hypothesis by using the aforementioned results from Trier et al. (2014) and assessing whether genomic cline shifts were more frequently in the direction of favoring the ancestral allele (i.e. the tree sparrow allele). For each locus, we converted the genomic cline parameter α to represent shifts in favor of the ancestral or derived allele, rather than one or the other parent species: negative values representing shifts towards the derived allele, and positive values shifts in favor of the ancestral allele. We then carried out intercept-only linear regression to test for a systematic deviation from α = 0, with a positive shift supporting our hypothesis. We also tested for significant skewness towards high α ancestry. These analyses were then repeated after filtering out 3 loci with tree sparrow minor allele frequency > 0.05, and again after filtering a further 7 loci with no pleiotropy data (see below), with no qualitative change in the results. Results for the full data set only are therefore presented.

### Genomic factors influencing ancestry reconstruction

To test which genomic factors influenced the degree of ancestry bias, we added predictor variables to the α ancestry regression analysis (above) and used model selection and model averaging (Burnham & Anderson 2002) in the R package MuMIn (Bartoń 2013). We hypothesized that nuclear-encoded proteins with a mitochondrial function (NEMPs) should be strongly biased in favor of the ancestral allele in Italian sparrows, particularly when the ancestral allele came from the same parent as the mitochondria (house sparrows). We identified NEMPs using the human MitoCarta database (Calvo et al. 2015). We created two binary variables: NEMP yes/no (6 loci) and house-ancestral NEMP yes/no (5 loci), which were never both included in the same model. More support for the latter variable would support our hypothesis. Pleiotropy may alter ancestry bias by changing the strength of selection and/or constraints on individual loci. To estimate pleiotropy, we counted the number of neighboring interacting proteins for each locus using the STRING database (http://www.string-db.org) for each of human, rat, mouse and chicken reference species, using the ‘get_neighbors’ function in the R package STRINGdb (Franceschini et al. 2013). Values from the different reference species were never included in the same model. We also included a binary variable for candidate hybrid-parent postzygotic incompatibility loci (henceforth PZIs) from Trier et al. (2014) and another to identify sex-linked loci. Pleiotropy values were logged prior to analysis and multiple regressions were run with 50 loci because 7 loci had no pleiotropy data for at least one reference species, and again with 47 loci after removing the 3 loci with tree sparrow minor allele frequency > 0.05, with no effect on the results. Results for 50 loci are presented. Furthermore, to examine whether different PZI categories differed in their degree of pleiotropy, we carried out ANOVA and post hoc Tukey Honestly Significant Difference (HSD) tests with chicken pleiotropy as the response (chicken was the best-fitting pleiotropy variable in the above regressions, see Results section). We used a single factorial predictor variable, with levels ‘neutral’, ‘hybrid-parent PZI’, ‘intraspecific incompatibility’, and ‘parental PZI’; the latter only including loci that were identified as PZIs in the parental house/Spanish sparrow genomic cline analysis (Hermansen et al. 2014) but were not in one of the previous two PZI categories, for this test.

### Hardy-Weinberg and linkage disequilibrium and unpurged genetic incompatibilities

Disequilibria within and between loci in a population can be caused by genetic drift, admixture between differentiated populations, or selection. With respect to selection caused by epistatic incompatibilities (DMIs), specific resulting patterns of disequilibria depend on the degree of dominance in the phenotypic expression of the ancestral allele, and on the symmetry of selection (for example whether selection is only against derived species 1/derived species 2 and no other allele combinations; Fig. S1). Using the Lago Salso population we compared evidence for the presence of DMIs or epistatic fitness effects more generally versus other sources of disequilibria (drift and admixture). As described below, we first tested for evidence of admixture, and then examined the distributions of Hardy-Weinberg disequilibria (HWD) and cross-chromosome linkage disequilibria (LD), and the genomic factors associated with variation in these values. Finally, we tested which locus pairs best fit a model of pairwise epistatic selection with dominance, and whether estimated selection coefficients matched the expectation for DMIs.

To test for any form of admixture, including through immigration from differentiated Italian sparrow populations, we first used the snmf function in the R package LEA (Frichot et al. 2015) to estimate k, the number of populations present in Lago Salso, with k = 1 representing no evidence of admixture. We tested k = 1:10, with each run initialized with all 57 loci. We calculated minimal cross-entropy across 50 repetitions for each value of k, using a proportion of 0.1 masked genotypes. We repeated this for values of the snmf parameter alpha of 1, 10, 100 and 1000, as this may influence results (Frichot et al. 2015). We also used the Bayesian assignment algorithm implemented in STRUCTURE (Pritchard et al. 2000). The correlated allele frequency model is often used in STRUCTURE analyses in order to identify subtle population structure. However, this was not our objective, and it is known that this model can create spurious structure and hence overestimate k (Pritchard et al. 2000). We therefore ran both correlated and uncorrelated allele frequency models for comparison. For each of k = 1:5, we ran both models 5 times, with 500k burnin followed by 1 million iterations. The optimal k was chosen using the Evanno method in Structure Harvester (Evanno et al. 2005; Earl & von Holdt 2012). All 57 loci and 86 individuals were used for both LEA and STRUCTURE.

We then estimated the distribution of ‘parental LD’ values (bias towards associations between alleles from the same parent species, called ‘ancestry LD’ by Schumer et al. 2014) in Lago Salso for 938 cross-chromosome locus pairs with minor allele frequency > 0.1 (46 loci). Cross-chromosome parental LD can be caused either by recent or ongoing gene exchange with the parent species (Barton 2000; Barton & Gale 1993; Gompert & Buerkle 2011; Fitzpatrick 2013) or segregating DMIs (Schumer et al. 2014; but see Schumer & Brandvain 2016). In order to factor out effects of inbreeding (Rogers & Huff 2009), we first calculated linkage disequilibrium, D, and then the correlation coefficient, r, as:

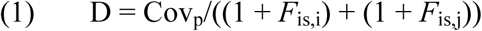

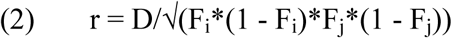

Where Cov_p_ = population (not sample) covariance of diploid genotypes scaled (0,1,2; 0 = house sparrow homozygote, 1 = heterozygote, 2 = Spanish sparrow homozygote), subscripts i and j represent the two loci, xs *F*
_is_ = inbreeding coefficient, and F = minor allele frequency. Positive r means positive associations between alleles from the same parent species. We used the distribution of r values to test for a bias towards positive parental LD, with a mean of zero indicating no bias. To examine the strength of LD in Lago Salso without reference to parental allele combinations, P values for r^2^ were calculated using equation 8 (T2 formula for unknown haplotype phase) from Zaykin et al. (2008)for two bi-allelic loci:

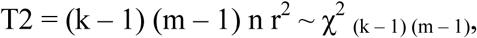

Where k and m indicate the number of alleles at each locus, and n is the number of individuals. To test for an overall significant r^2^ across all locus pairs, the difference between the actual mean p value and the mean of 1000 permuted (diploid genotypes permuted among individuals for each locus) data sets was calculated. Threshold-specific false discovery rate (FDR) was also calculated, at 100 p-value thresholds from 0.001 to 0.1, as mean ((N permuted locus pairs below threshold)/(actual N locus pairs below threshold)). The number of true positives at each p value threshold was calculated as (actual N locus pairs below threshold)-mean(N permuted locus pairs below threshold). (see example code in supplemental data for full description). Furthermore, mean r^2^ per locus was used in multiple regression model selection to examine the impacts of genomic architecture on LD. We included the following predictor variables: sex linkage, internal incompatibilities, parental PZIs, parent of origin of ancestral allele, pleiotropy (number of neighboring proteins in chicken), parental average minor allele frequency, and difference in allele frequency between parents. The latter two may differ when the same allele is the minor allele in both parents. Hybrid-parent PZIs were all excluded from the analysis due to low minor allele frequency in Lago Salso (<=0.1). We also tested for HWD at individual loci and combined significance across all loci using the least squares based method in Genodive (Meirmans & Van Tienderen 2004), and carried out the same genomic architecture regression analyses on resulting *F*_is_, and absolute *F*_is_, values as for LD.

Deviations from HWE and LE combined can provide information on the pattern of selection acting on a locus pair (e.g. Fig. S1). We used this information by fitting a model of epistatic viability selection and dominance to the full cross-chromosome pairwise genotype data (938 locus pairs). We assumed that the current generation was at HWE and cross-chromosome LE prior to viability selection, and estimated by maximum likelihood the ancestral allele frequency at each locus prior to selection, the dominance of the ancestral over the derived allele (both one parameter per locus) and, for each locus pair, the estimated coefficient of selection against four different allelic combinations: house_i_/house_j_, Spanish_i_/Spanish_j_, house_i_/Spanish_j_ and Spanish_i_/house_j_ (i and j represent the first and second locus in a pair), taking into account which parent species provided the ancestral allele for each locus. The general ancestral/derived formulae for the nine pairwise diploid genotypes, not accounting for parent of origin, was:

F(AA_i_AA_j_) = E(AA_i_AA_j_)-s(A_i_A_j_) E(AA_i_AA_j_)
F(Ad_i_AA_j_) = E(Ad_i_AA_j_)-s(A_i_A_j_) E(Ad_i_AA_j_) D_i_-s(d_i_A_j_) E(Ad_i_AA_j_) (1-D_i_)
F(dd_i_AA_j_) = E(dd_i_AA_j_)-s(d_i_A_j_) E(dd_i_AA_j_)
F(AA_i_Ad_j_) = E(AA_i_Ad_j_)-s(A_i_A_j_) E(AA_i_Ad_j_) D_j_-s(A_i_d_j_) E(AA_i_Ad_j_) (1 – D_j_)
F(Ad_i_Ad_j_)=E(Ad_i_Ad_j_)-s(A_i_A_j_) E(Ad_i_Ad_j_) D_i_ D_j_-s(A_i_d_i_) E(Ad_i_Ad_j_) D_i_ (1-D_j_)-s(d_i_A_j_) E(Ad_i_Ad_j_) (1-D_i_) D_j_-s(d_i_d_j_) E(Ad_i_Ad_j_) (1-D_i_) (1-D_j_)
F(dd_i_Ad_j_) = E(dd_i_Ad_j_)-s(d_i_A_j_) E(dd_i_Ad_j_) D_j_-s(d_i_d_j_) E(dd_i_Ad_j_) (1-D_j_)
F(AA_i_dd_j_) = E(AA_i_dd_j_)-s(A_i_d_j_) E(AA_i_dd_j_)
F(Ad_i_dd_j_) = E(Ad_i_dd_j_)-s(A_i_d_j_) E(Ad_i_dd_j_) D_i_-s(d_i_d_j_) E(Ad_i_dd_j_) (1-D_i_)
F(dd_i_dd_j_) = E(dd_i_dd_j_)-s(d_i_d_j_) E(dd_i_dd_j_)

Where A and d indicate ancestral and derived alleles respectively, s = the four selection coefficient parameters (range 0-1), D = ancestral allele dominance parameter (range 0-1; 0.5 = additivity), F = post-selection genotype frequency, and E = genotype frequency at HWE and LE prior to selection, given the parameter value for prior allele frequency at each locus. Parent of origin of the ancestral allele was accounted for by altering the incorporation of dominance. Post-selection frequencies were then scaled to proportions before fitting to the data using a multinomial model (see example code in supplemental data for full description). In the maximum likelihood model all parameters (four pairwise selection coefficients per locus pair, and per-locus ancestral allele dominance and prior allele frequencies) were updated simultaneously based on their individual likelihoods each iteration, using a Metropolis algorithm (Gelatt & Vecchi 1983). After extensive testing, we chose a MCMC strategy of 10 random sets of starting values for all parameters, each followed by 100k MCMC iterations. The maximum of the summed likelihoods across all locus pairs was chosen as the best model. This model does not represent a simulation of selection on a true population, but rather quantifies the fit of each locus pair to the global ML set of parameter values, given a single value for prior allele frequency and ancestral dominance per locus applied to all locus pairs involving that locus. It hence provides a ranked quantification of the fit of each locus pair to the model of pairwise epistatic selection. The estimated selection coefficients were then used to examine the extent to which well-fitting locus pairs followed the expectations of DMIs (symmetric selection against heterospecific genotypes, or selection against heterospecific derived-derived combinations only).

## RESULTS

### Deviation towards the ancestral allele

The ancestral alleles identified through tree sparrow genotyping were evenly distributed among the two parent species (28 loci with higher ancestral allele frequency in the Spanish sparrow, and 29 higher in house sparrow; Table S1). There was a significant bias in α ancestry towards the ancestral allele across all loci (intercept=0.25, SE=0.11, d.f. = 56, t=2.31, *P*=0.025; Fig. 2). The distribution of these cline centers was also significantly skewed towards ancestral alleles (skewness: 1.07 standard error: 0.32; Z_skewness_=3.37), supporting a general trend of fixation of the ancestral allele through selection.

**Fig. 1.**
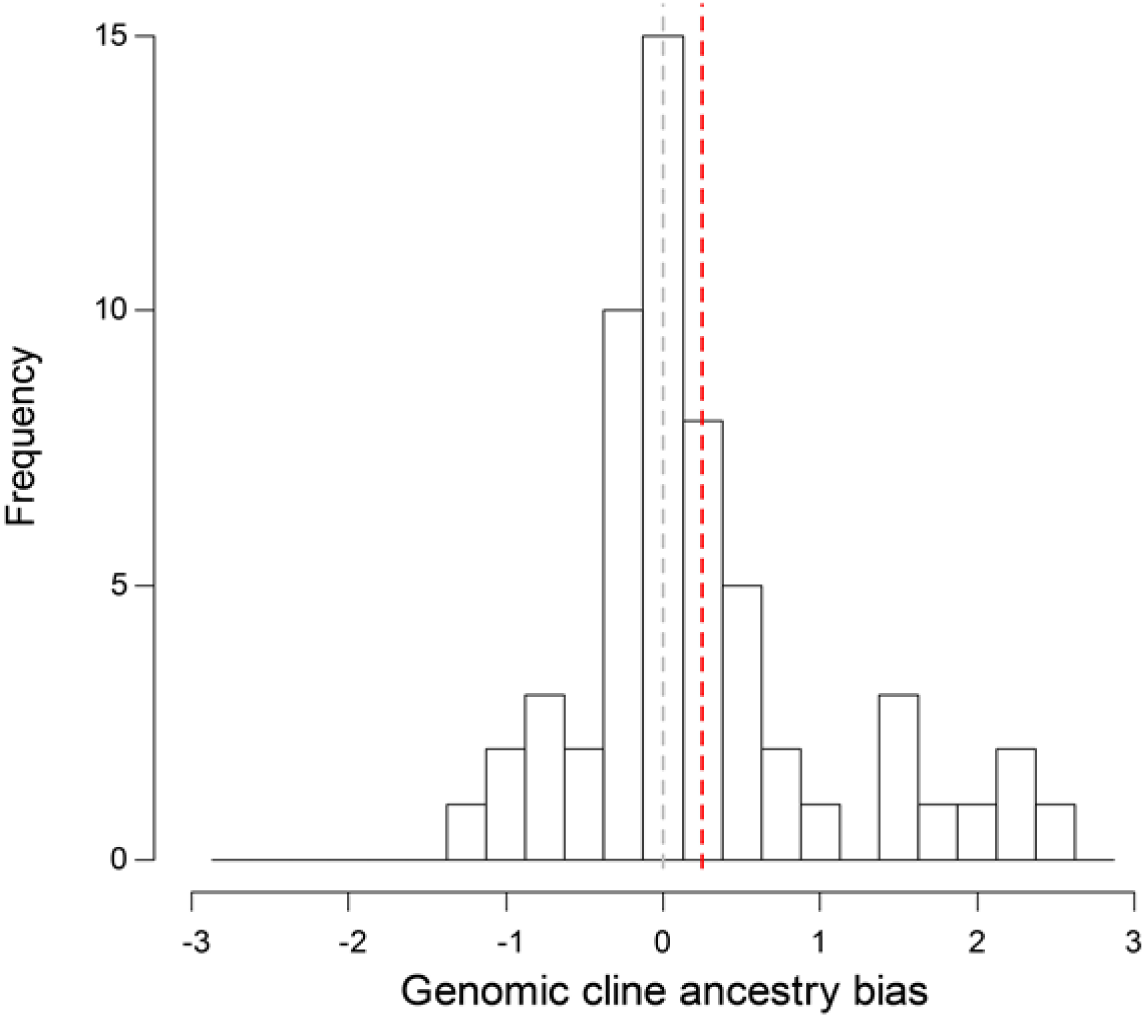
Distribution of the ancestral genomic cline center (α ancestry) among 57 SNPs. Positive values indicate bias in favor of the ancestral allele. Grey dashed vertical line = 0; red dashed vertical line = mean α ancestry.

**Fig. 2.**
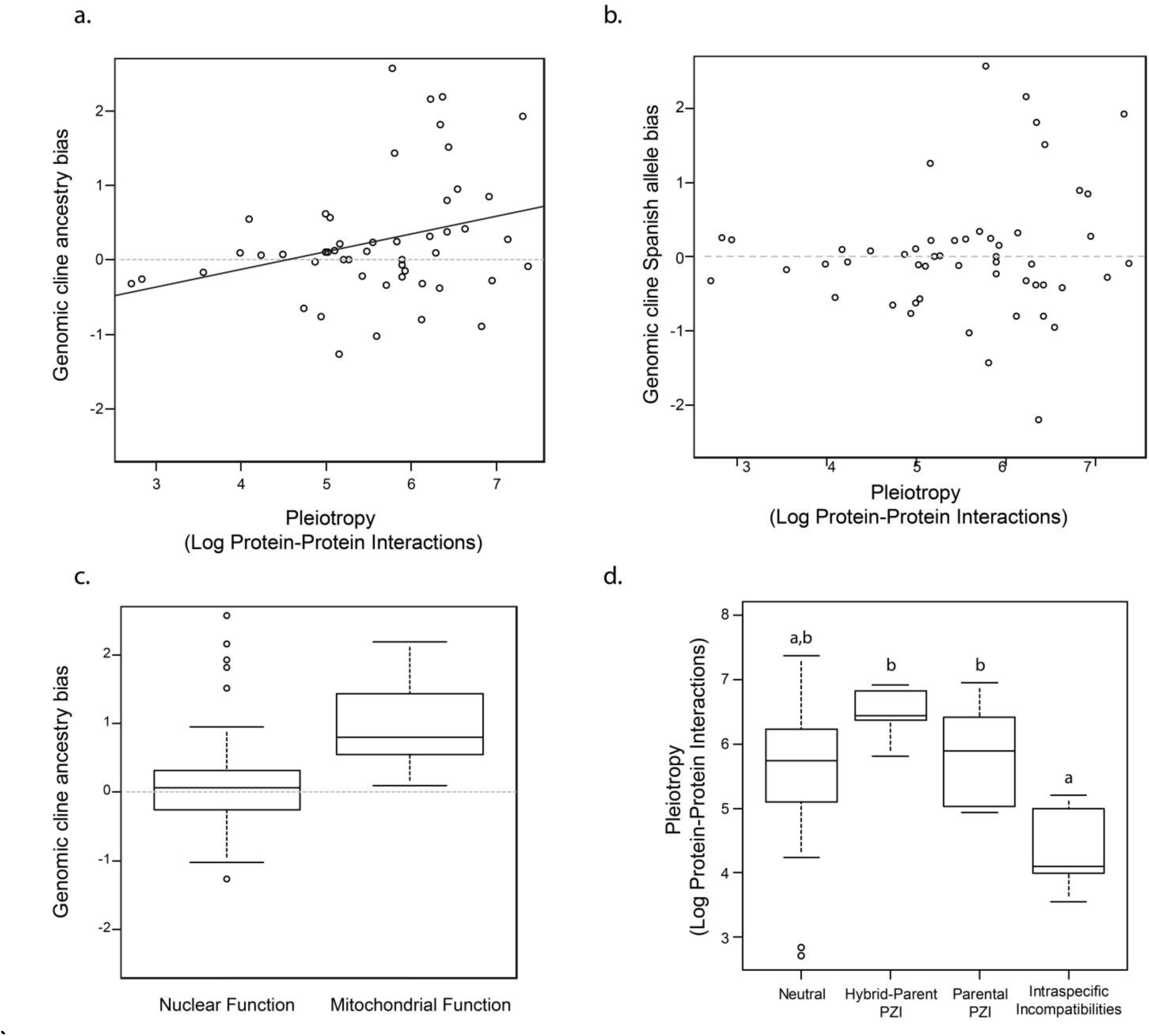
Effect of genomic properties on genomic cline α. Pleiotropy (number of neighboring interacting proteins in chicken) against (a) α ancestry and (b) α for bias in favor of alleles origination from Spanish sparrows. (c) Effect of mitochondrial function (house-ancestral NEMPs). (d) Variation in pleiotropy depending on which type of PZI the loci investigated are; letters indicate post hoc groupings.

Multiple regression and model averaging on α ancestry revealed that pleiotropy was the most important predictor variable, with increased pleiotropy loci leading to significantly more bias in favor of the ancestral allele (Table 1, Fig. 3a,b). House-ancestral NEMPs were the second most important predictor, being biased in favor of the ancestral allele (Figure 3c). Adding the single Spanish-ancestral NEMP reduced significance, supporting the hypothesis that only derived NEMP alleles originating from Spanish sparrows were selected against. Sex-linked loci and hybrid-parent PZIs were not significantly more biased in favor of the ancestral allele than the rest. We also found that PZI categories differed significantly in pleiotropy (one way ANOVA, 50 loci: df=3.46, F = 4.4, *P*=0.008; 47 loci: df=3.43, F=4.15, *P*=0.011). Incompatibilities segregating within Italian sparrows (internal incompatibilities) had lowest pleiotropy, significantly lower than hybrid-parent PZIs (post hoc test, *P* =0.006 for 50 loci; *P*=0.007 for 47 loci), which had the highest mean pleiotropy (Figure 3d). Intraspecific incompatibilities also had marginally significantly lower pleiotropy than parental PZIs (*P*=0.031; reduced to *P*=0.052 with 47 loci) and marginally non-significantly lower than neutral loci (*P*=0.053; *P*=0.065 with 47 loci).

**Table 1.** Multiple regression model averaging for predictors of ancestrybias (genomic cline α ancestry). ^1^Sum of Akaike weights over all models including the explanatory variable (Burnham and Anderson 2002).

| Variable | Importance<sup>1</sup> | Estimate<br>(SE) | z<br>value | p value |
| --- | --- | --- | --- | --- |
| (Intercept) | NA | -0.95<br>(0.78) | 1.21 | 0.23 |
| Pleiotropy | 0.81 | 0.25<br>(0.11) | 2.23 | **0.03\*** |
| House-ancestral<br>mitochondrion | 0.63 | 0.89<br>(0.40) | 2.21 | **0.03\*** |
| PZI | 0.39 | 0.51<br>(0.42) | 1.17 | 0.24 |
| Mitochondrion | 0.37 | 0.72<br>(0.36) | 1.94 | **0.05.** |
| Sex-linked | 0.32 | -0.23<br>(0.25) | 0.90 | 0.37 |

**Fig. 3.**
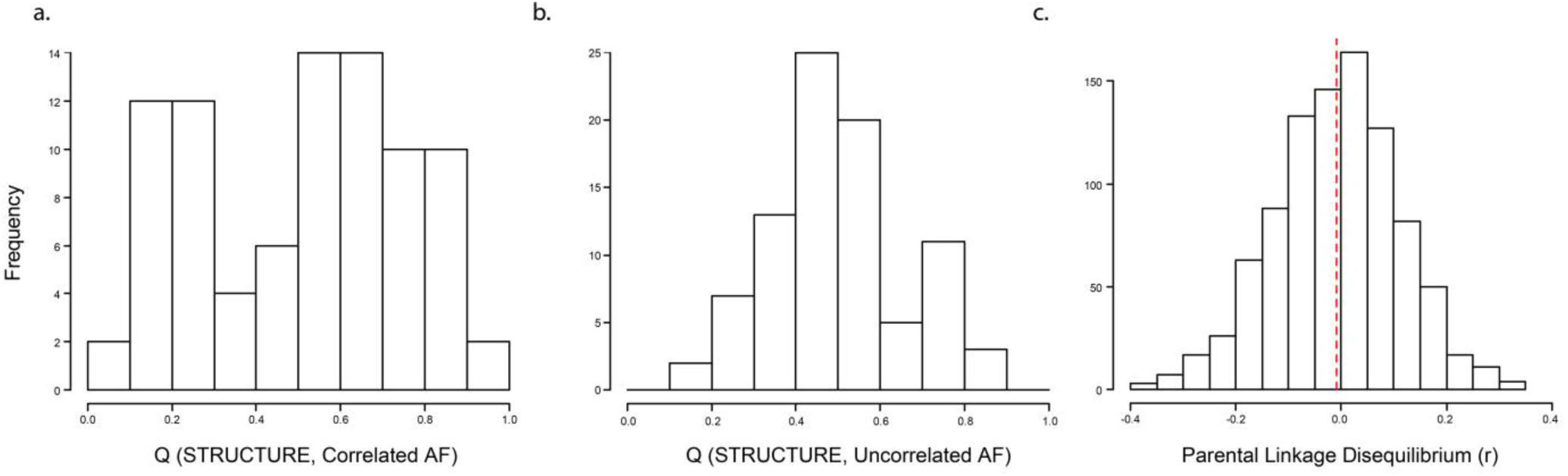
Population structure and admixture in Lago Salso. (a) STRUCTURE Q value histogram for the correlated allele frequency model. (b) Q values for the uncorrelated allele frequency model. (c) The distribution of parental LD correlation coefficients among 938 cross-chromosome locus pairs (red dashed vertical line = mean r).

### Unpurged Dobzhansky-Muller incompatibilities in Lago Salso

LEA analyses uniformly supported the presence of a single population in Lago Salso (Figure S2), hence indicating no recent admixture between Lago Salso Italian sparrows and other, differentiated populations of Italian sparrows or either parent species. However, both correlated (CAF) and uncorrelated allele frequency (UAF) STRUCTURE models supported k=2 (Figure 4a,b; Supplemental data). The histogram of Q values (probability of being a member of one of the two clusters) for the uncorrelated allele frequency model is unimodal, with a single peak at Q=0.5. This is not to be expected in cases of admixture, and hence the population structure is more likely to be caused by drift or epistasis linked to the hybrid properties of this species. Furthermore, there was no evidence of an excess of cross-chromosome parental LD in this population, with the mean pairwise parental correlation coefficient very close to zero and slightly negative (Figure 4c). Therefore we found no strong evidence for either ongoing admixture with the parent species, or an excess of cross-chromosome parental genotypes caused by pervasive segregating incompatibilities (cf. Schumer et al. 2014; Schumer & Brandvain 2016).

**Fig. 4.**
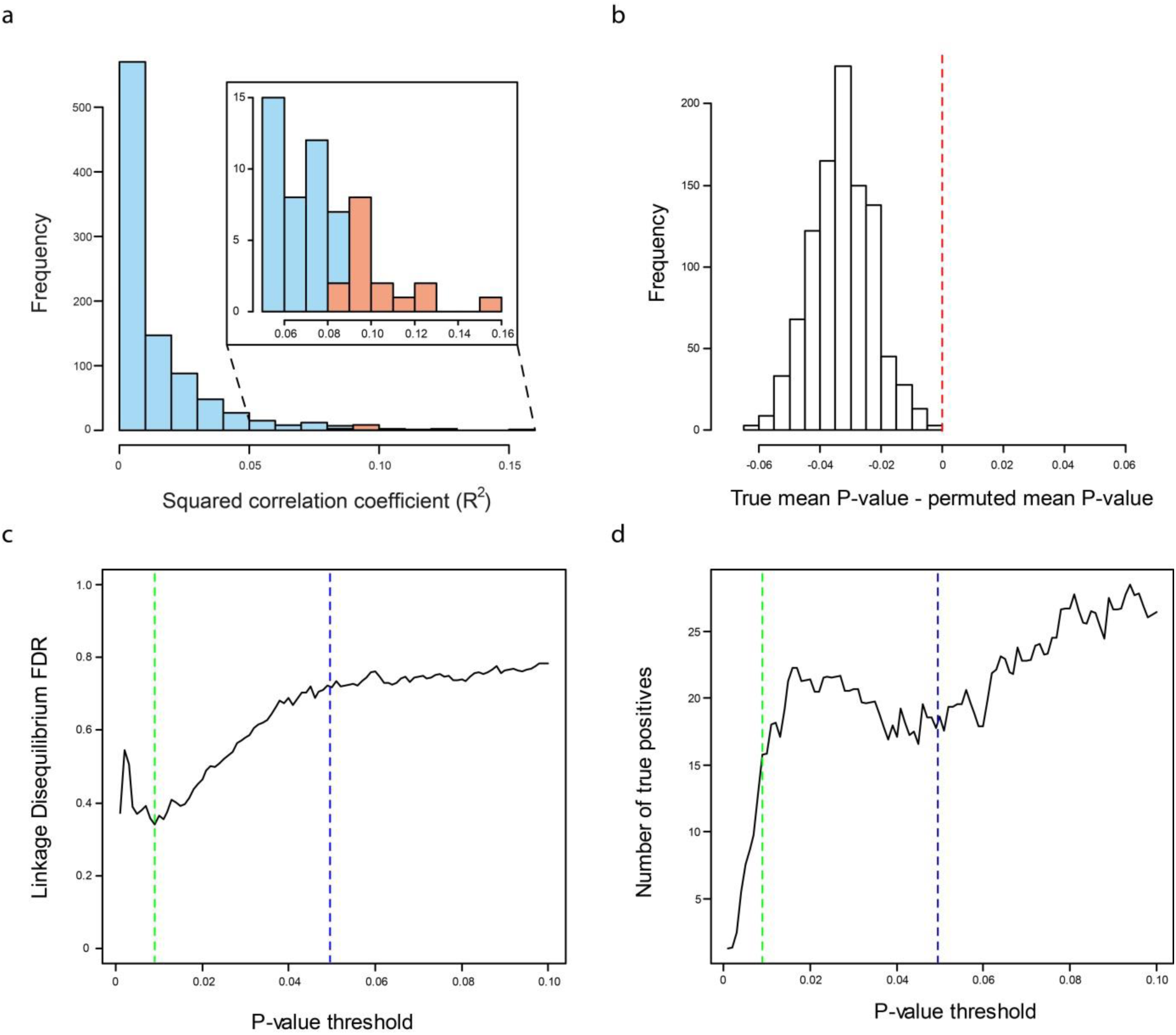
Linkage disequilibria in the Lago Salso population. (a) Histogram of r^2^ values. The 16 true significant locus pairs at p = 0.009 are shown in red. (b) The difference between true mean and permuted mean r^2^. Absence of overlap with 0 (red dashed vertical line) over 1000 permutations indicates significant overall LD. (c) False discovery rate (FDR) for significantpairwise LD at different p value thresholds. Green dashed vertical line: minimum FDR of 34% at p = 0.009; blue dashed line: P value threshold for 5% false discoveries across all tests (47/938 tests; p = 0.0495). (d) Estimated number of true positive pairs in LD. Green and blue lines as in panel c.

However, our results suggest there is persistent cross-chromosome linkage disequilibrium in the Lago Salso population (Figure 5a). The mean r^2^ of 0.014 was significantly higher than expected by chance (Figure 5b). The minimum threshold FDR was quite high, being 34% at *P* =0.009. The estimated number of true positives at *P* =0.009 was 16 (Figure 5a; c-d). While these disequilibria might be caused by drift, we would not expect associations with genomic architecture under that scenario. Furthermore, we found that loci classified as parent-parent PZIs (excluding internal Italian or hybrid-parent PZIs) had significantly increased mean r^2^ (linear regression: df=1,44; t=2.4; *P*=0.02; Figure 6a). In addition, r^2^ increased significantly with decreasing mean parental minor allele frequency (df =1.44; t=-2.16; *P*=0.04; Figure 6b), indicating that loci closer to fixation in the parents were more likely to be involved in epistatic interactions in the hybrid. However, these two variables were non-significant in a multiple regression and not significant at *P*=0.05 using model averaging (parental PZI *P*=0.07; parental minor allele frequency *P*=0.18), and hence further verification of these effects is required. Two of 57 loci were in significant heterozygote deficit and one in significant excess (Table S1). Across all loci, the Lago Salso population was found to be in significant heterozygote deficit (*F*_is_=0.03, *P*=0.028). In the best linear regression model according to AICc on *F*_is_, internal incompatibility loci and parental allele frequency difference were marginally non-significant (model linear regression: df=2,43; F=2.4; *P*=0.1): internal incompatibilities had higher heterozygote deficit (*P* =0.07).Loci with higher parental allele frequency difference tended towards heterozygote excess. For absolute *F*_is_, the best model with the lowest AICc was significant (df=3.42; F=3.6; *P*=0.02) and showed that parental PZIs had stronger HWD than the rest (*P*=0.003), while sex-linked loci had reduced HWD (*P* =0.05), and higher parental mean minor allele frequency non-significantly increased HWD. Using model averaging, Parental PZIs remained significant (*P*=0.01), while sex linkage was marginally significant (*P*=0.06).

**Fig. 5.**
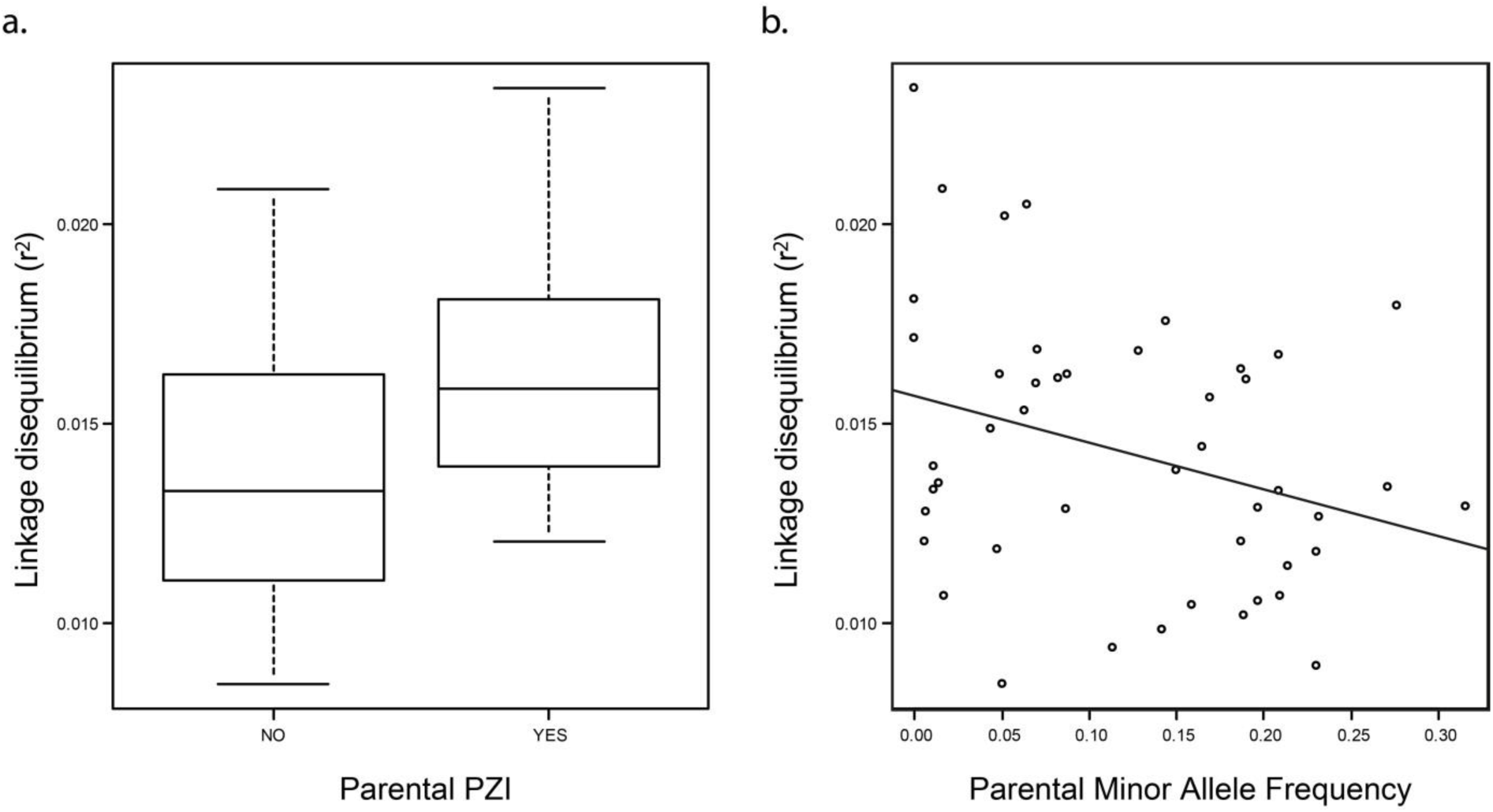
Genome-level factors influencing linkage disequilibrium in Lago Salso. (a) Boxplot of linkage disequilibrium (r^2^) for parental PZI loci or other loci. (b) Regression between linkage disequilibrium (r^2^) against parental minor allele frequency.

In the epistatic selection ML analyses, the strongest evidence for cross-chromosome pairwise epistatic selection among the 46 tested loci was between GSTK1 (chromosome 1) and HECTD1 (chromosome 5) (Table S2). This pair was also in strongest LD, and HECTD1 had the highest heterozygote deficit of all loci (Table S1). However, the strongest selection coefficient was not against heterospecific allele combinations but against Spanish/Spanish allele combinations (s=0.81) and the weakest against derived Spanish/derived house combinations (s=0.004), contrary to expectations. Among the best-fitting locus pairs, with maximum likelihood improvement over the null model > 4 units, there was no bias in selection coefficients against heterospecific allele combinations *per se*, or against heterospecific derived/derived combinations (Table S2), and hence consistent with the absence of a bias in favor of parental LD.

## DISCUSSION

Genetic incompatibilities are widespread (Crespi and Nosil 2013) and may have severe fitness consequences in admixed populations (Corbett-Detig 2013), including hybrid species. We find support for the hypothesis that genetic incompatibilities have shaped genome evolution in the Italian sparrow, and continue to do so. First, our data support the hypothesis that compatible ancestral allele combinations have been recreated in the hybrid genome, disfavoring derived alleles likely to be present in the same individual for the first time in the hybrid taxon. This selection probably occurred during the process of sorting of incompatibilities that led to reproductive isolation between the two parent species and the emerging hybrid lineage (Hermansen et al. 2014). As predicted, nuclear loci with a mitochondrial function exhibit a strong bias in favor of ancestral alleles in the hybrid species, particularly when the ancestral allele is inherited from the same parent as the mitochondria (house sparrow), indicating selection against derived alleles that have not previously interacted with house sparrow mitochondria. This pattern of ancestry bias extends to loci not previously identified as candidate incompatibility loci. We also found that loci with higher pleiotropy are more shifted towards ancestry, supporting the hypothesis that higher pleiotropy leads to stronger directional selection for purging of incompatibilities in hybrids. Interestingly, we found that candidate incompatibility loci still segregating within Italian sparrows had very low pleiotropy values. In addition, we found several lines of evidence consistent with pervasive ongoing epistatic fitness interactions among loci within one population, particularly involving loci previously identified through genomic cline analysis as incompatibilities between the parents, but not between hybrid and parent or within the hybrid taxon. Many of these interactions did not appear to represent classic DMIs, either as symmetric selection against mixed genotypes or selection against heterospecific derived/derived combinations only.

Our findings are, to the best of our knowledge, the first empirical evidence for the predicted bias towards ancestral genotype reconstruction in a hybrid lineage (Gavrilets 1997; Shuker et al. 2005) during the formation of a stabilized and viable hybrid genome (Rieseberg et al. 1995). This suggests that sorting of parental incompatibilities (Hermansen et al. 2014) may be a quite deterministic process. The pervasiveness of this phenomenon - with ancestral alleles seemingly favored for many more loci than those previously identified as incompatibility loci-suggests that many loci in the hybrid genome harbor a potential for conflict between alleles from the different parent species, without playing any well-defined role in terms of post-zygotic isolation between the hybrid species and its parents. In many circumstances ancestral alleles might be expected to form weak hybrid-parent barriers as they should be cross-compatible with derived alleles with which they have previously coexisted (Schumer et al. 2015), and some incompatibility loci may have more moderate effects on fitness, which would not necessarily lead to sterility or mortality as often assumed under the DMI model (Fang et al. 2012; Schumer et al. 2014). We therefore suggest – consistent with our results - that more pleiotropic loci are more likely to form strong hybrid-parent barriers. This may be because the ancestral allele from one parent species at a particular locus in the hybrid taxon is likely to interact with a greater number of derived alleles from other parent species at other loci.

Incompatibilities involving mitochondrial DNA and nuclear genes with mitochondrial functions (NEMPs) are thought to be common (Burton and Barreto 2012). Italian sparrows inherited their mitochondria from the house sparrow (Hermansen et al. 2011; Elgvin et al. 2011; Trier et al. 2014), and mtDNA forms a strong reproductive barrier at the boundary with Spanish sparrows (Trier et al. 2014). Purging of incompatible alleles at individual loci within the mitochondria is an inherently slow process due to the lack of mitochondrial recombination. We hence propose that the high propensity towards fixation of ancestral NEMP alleles from the house sparrow may be due to strong selection against derived Spanish alleles, incompatible with the potentially numerous derived alleles in the house sparrow mitochondrial genome. As predicted, since there is no expectation for being incompatible with the house sparrow mitochondrial genome, the effect was smaller when the single NEMP locus for which the derived allele was inherited from the house sparrow was included. Together, these findings support the hypothesis that in the Italian sparrow, the mitochondrial genome constitutes an important source of past and present inter-genomic conflicts, likely involving metabolic pathways (Trier et al. 2014).

Evidence that pleiotropy has major effects on gene evolution and expression, slowing down divergent directional selection and constraining variation in gene expression, is mounting (Mank et al. 2008; Papakostas et al. 2014; Uebbing et al. 2016). Here we present the first evidence that pleiotropy also has important impacts on hybrid genome evolution. Given the high pleiotropy of strongly ancestry-shifted loci and the disproportionate role of mito-nuclear interactions, we speculate that loci interacting with large numbers of differentiated loci have a strong and deterministic influence on hybrid genome evolution, favoring ancestral alleles from the same parent species. Particularly intriguing is the pattern of exceptionally low pleiotropy among loci previously identified as candidate incompatibilities segregating within Italian sparrows, with their steep but only weakly shifted genomic clines. Low pleiotropy may be required for incompatibilities segregating within a taxon to evolve independently from the rest of the genome, and hence develop relatively narrow genomic clines.

The strong overall heterozygote deficit (as opposed to random deviations from HWE) in the focal Lago Salso population could be caused by population subdivision, which we regard as unlikely given our population structure results, but cannot be predicted by drift. However, we suggest that if there is pervasive epistatic selection among the loci studied here, regardless of whether it is selection linked to DMIs, this should lead on average to positive *F*_is_. This is because selection against dominant/dominant allele combinations causes positive *F*_is_ and is also more effective than selection on other combinations of dominance levels, as it acts in all heterozygote genotypes. This more effective dominant/dominant selection should lead to a disproportionate effect of these locus combinations on HWD, and hence to an average heterozygote deficit. Weaker HWD on the Z chromosome than autosomes may be due to its hemizygous nature, leading to incompatibilities and recessive deleterious alleles being purged faster (Charlesworth et al. 1987; Borge et al. 2005; Ellegren 2009; Trier et al. 2014), and hence leaving little segregating variation present within the hybrid species at loci with strong epistatic fitness effects.

The best evidence that the high linkage disequilibria and heterozygote deficit in the Lago Salso population are caused by selection rather than admixture comes from the significant associations between disequilibria and aspects of genomic architecture related to conflicts between parental genomes. Loci previously identified as candidate incompatibilities isolating the parent species, but not isolating the hybrid from its parents, show both strong HWD and strong mean LD, suggesting that these loci may be more constrained in their evolution within the hybrid species than other categories, still having important fitness effects despite not forming narrow clines. Given that we found no strong evidence that epistatic fitness effects represented classic DMIs in Lago Salso, it is unclear yet whether this hybrid species differs in the pervasiveness of epistasis from non-hybrid species (Corbett-Detig et al. 2013). Similar tests of associations between disequilibria and genomic architecture in other systems would be useful. On the other hand, the apparent importance of candidate incompatibilities in this population suggests that fitness effects may nevertheless differ in hybrids due to parental differentiation, albeit specific fitness effects may not always fit the expectations of DMIs. The strongest support of this hypothesis was for an interaction between HECTD1 and GSTK1; the former being a candidate parental (but not hybrid-parent or internal Italian) incompatibility and the latter a candidate internal Italian sparrow incompatibility. We found no existing evidence of known interactions between these genes in other taxa.

Distinguishing selection from other forces such as drift and admixture as the cause of disequilibria, or of steep or shifted genomic clines, remains challenging. At this point for example, we cannot entirely exclude a role for assortative mating or population subdivision in generating disequilibria in Italian sparrows at the population level. However, we highlight that non-random associations between cline parameters or disequilibria and genome-level variables such as pleiotropy and ancestry may provide evidence for selection. Such tests could in the future complement other statistical methods being developed for natural admixed populations, alongside manipulative experiments.

The potentially creative role of hybridization in evolution is currently much discussed (e.g. Abbott et al. 2013; Seehausen et al. 2014). Hybridization may enhance evolvability due to increased genetic variation (Barton 2001) and induce evolutionary novelty through transgressive segregation (Rieseberg et al. 1999). Hybrid speciation is in itself a good example of the creative role that hybridization may play in evolution (Mallet 2007; Abbott et al. 2013). Although hybrid species have been found to readily adapt (Rieseberg et al. 2003; Heliconius Genome Consortium 2012; Eroukhmanoff et al. 2013b), admixed genomes may inherit incompatibilities that severely reduce their viability and restrict evolvability to a few limited directions in genotype space. Here, we find evidence suggesting that hybrid speciation can have lasting impacts on genetic architecture. Epistatic interactions among divergent loci can persist within a hybrid species and may reduce fitness long after hybridization initially occurred and hybrid-parent reproductive isolation has evolved. Further work is needed to examine the extent to which such segregating loci in Italian sparrows are facilitating divergence and adaptation or hampering evolution. A larger sequencing effort across the entire genome and the inclusion of additional outgroup species, combined with more experimental work on laboratory-generated hybrids (e.g. Eroukhmanoff et al. 2016), would likely shed more light on this phenomenon. More work is thus needed to unravel the complex effects hybridization may have on organismal diversity, especially in the case of hybrid speciation.

## Acknowledgements

We thank S. A. Sæther and numerous assistants for their help during the field work, B. Dogan for help with laboratory work, and K. Voje for helpful comments on a previous draft of the manuscript. This work was supported by The Research Council of Norway, Molecular Life Science (MLS), University of Oslo, Centre for Ecological and Evolutionary Synthesis (CEES), University of Oslo, The Swedish Research Council, The Wenner-Gren Foundation and The Marie Curie Foundation. We declare no conflict of interest. Data will be archived in DRYAD upon acceptance.

## Supporting information

**Fig. S1.**
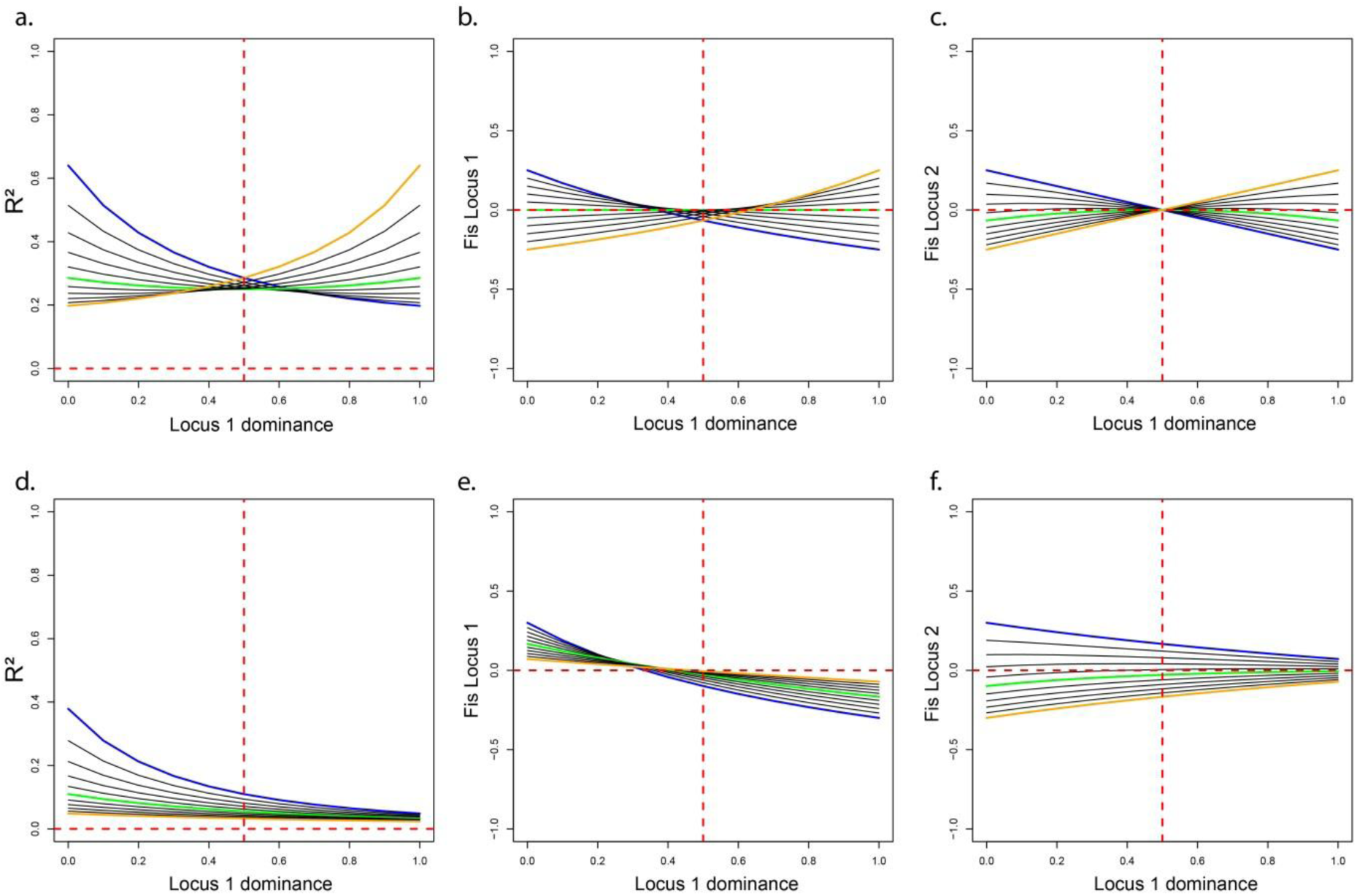
The effects of dominance on linkage and Hardy-Weinberg disequilibrium, caused by within-generation viability selection against heterospecific two-locus genotypes. a-c: symmetric selection against heterospecific genotypes. d-f: asymmetric selection against derived species A/derived species B allele combinations only. Dominance refers to the ancestral allele; additivity = 0.5. Green line = additivity locus 2; blue line = fully recessive ancestral allele locus 2; orange line = fully dominant ancestral allele locus 2 (black lines are intervening dominance values). Horizontal dashed red lines indicate LE or HWE; vertical dashed red lines indicate additivity of locus 1. Selection = 1 in all cases. In this example, heterospecific genotypes are either derived-derived or ancestral-ancestral.

**Fig. S2.**
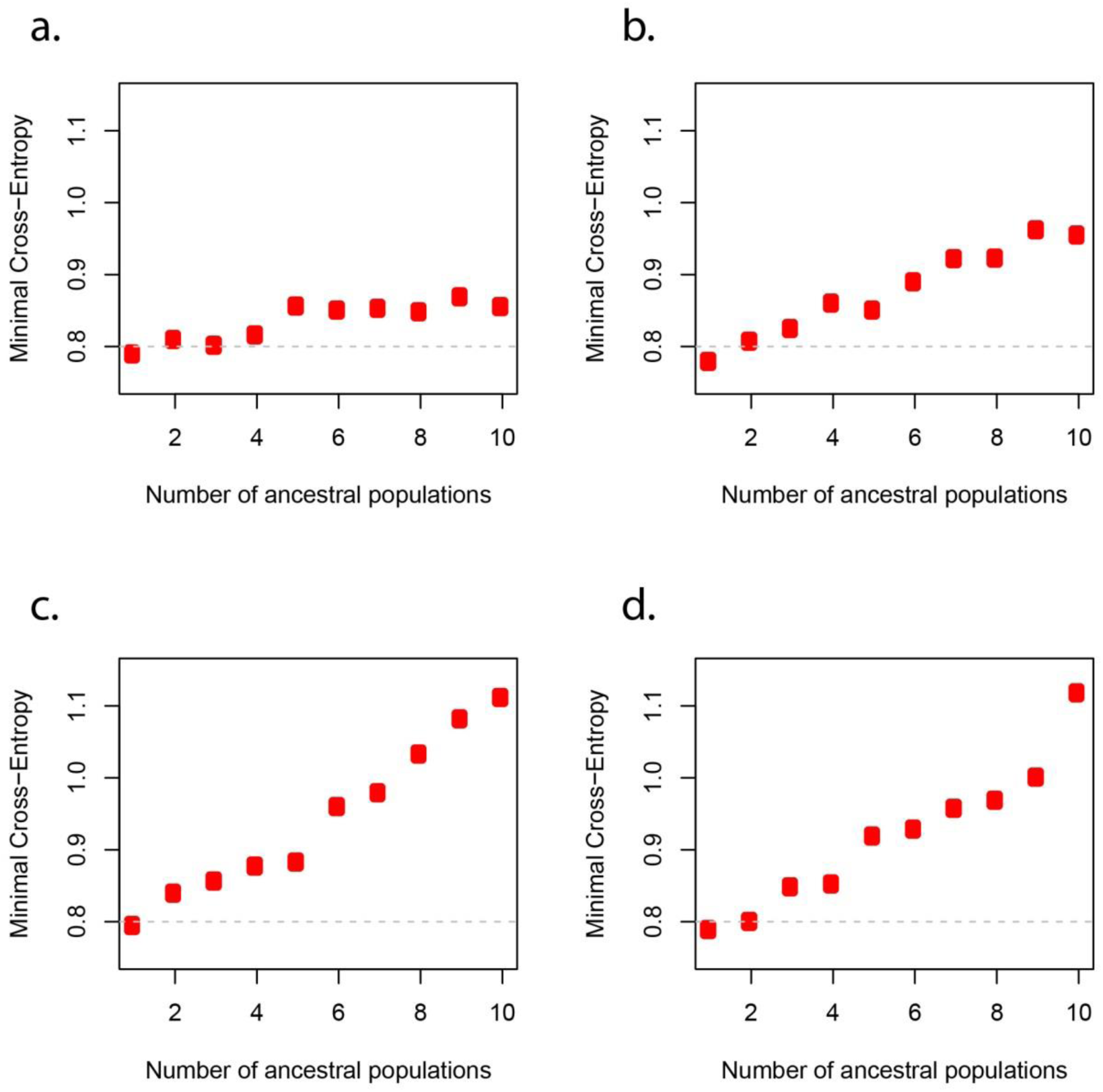
Minimal cross-entropy for each of k (number of clusters) = 1-10, for snmf parameter alpha = (a) 1, (b) 10, (c) 100, (d) 1000. The horizontal grey dashed line is an arbitrary reference for comparison between panels. Lowest minimal cross-entropy indicates best fit.

